# Hydrogels with stiffness-degradation spatial patterns control anisotropic 3D cell response

**DOI:** 10.1101/2023.01.25.525504

**Authors:** Claudia A. Garrido, Daniela S. Garske, Mario Thiele, Shahrouz Amini, Samik Real, Georg N. Duda, Katharina Schmidt-Bleek, Amaia Cipitria

## Abstract

In nature, tissues are patterned, but most biomaterials used in human applications are not. Patterned biomaterials offer the opportunity to mimic spatially segregating biophysical and biochemical properties found in nature. Engineering such properties allows to study cell-matrix interactions in anisotropic matrices in great detail. Here, we developed alginate-based hydrogels with patterns in stiffness and degradation, composed of distinct areas of soft non-degradable (Soft-NoDeg) and stiff degradable (Stiff-Deg) material properties. The hydrogels exhibit emerging patterns in stiffness and degradability over time, taking advantage of dual Diels-Alder covalent crosslinking and UV-mediated peptide crosslinking. The materials were mechanically characterized using rheology for single-phase and surface micro-indentation for patterned materials. 3D encapsulated mouse embryonic fibroblasts (MEFs) allowed to characterize the anisotropic cell-matrix interaction in terms of cell morphology by employing a novel image-based quantification tool. Live/dead staining showed no differences in cell viability but distinct patterns in proliferation, with higher cell number in Stiff-Deg materials at day 14. Patterns of projected cell area became visible already at day 1, with larger values in Soft-NoDeg materials. This was inverted at day 14, when larger projected cell areas were identified in Stiff-Deg. This shift was accompanied by a significant decrease in cell circularity in Stiff-Deg. The control of anisotropic cell morphology by the material patterns was also confirmed by a significant increase in filopodia number and length in Stiff-Deg materials. The novel image-based quantification tool was useful to spatially visualize and quantify the anisotropic cell response in 3D hydrogels with stiffness-degradation spatial patterns. Our results show that patterning of stiffness and degradability allows to control cell anisotropic response in 3D and can be quantified by image-based strategies. This allows a deeper understanding of cell-matrix interactions in a multicomponent material.

## 1 Introduction

Patterns are naturally occurring in nature, macroscopically and microscopically. The constant remodeling of the extracellular matrix (ECM) leads to emergent patterns of cells, ECM properties and cell behavior (1)(2). Biomaterials like hydrogels are a useful tool to study cell-matrix interaction as they can mimic various characteristics of the cell niche (3). Multiple approaches have been taken to study cell response to specific ECM properties, for example: materials with different stiffness to study focal adhesions (4) and mechanosensation (5), stress relaxing materials to mimic the viscoelastic behavior of biological tissues (6), independent control of mechanical properties and fibronectin presentation for stem cell engineering (7), modifications in the scaffold architecture and pore distribution (8), or biomolecule presenting/releasing materials (9). Patterned materials will offer the opportunity of imitating and guiding cell behavior with a closer relation to the natural counterpart.

Alginate is natural, biocompatible and inert polymer. Its versatile structure allows modifications to modulate key biophysical cues. Chemical modifications of the alginate structure, such as thiolation (10), oxidation (11), amidation (12) and Diels-Alder addition (13)(14) can be the base to implement additional crosslinking, improve or control degradation behavior or enable a controlled drug release. Alginate is capable to be crosslinked by various means such as ionic and covalent crosslinking (15). That capability opens the possibility to mimic and control distinct ECM properties. Alginate can thus be made such that a relatively broad range of mechanical properties can be covered or a dynamic environment provided to cells (11)(16).

Multiple biophysical and biochemical factors contribute to the complexity of the ECM. The interplay between these factors is a current topic of research. The mechanical properties of the ECM have been examined in single-phase 3D hydrogels with different elastic modulus, showing that the stiffness has an effect on cell phenotype (17)(18) and cell migration (19). The degradability of the material is important to create dynamic 3D matrices and it can affect cell spreading, cell interactions (20) and morphology (21)(22). Fewer studies investigate the interaction of stiffness and degradation on cell behavior in 3D encapsulated cells. Previous research showed that the simultaneous modulation of stiffness and degradation can influence cell proliferation or differentiation (23) and thereby control cell phenotypes (24).

The combination and spatial patterning of biophysical and biochemical cues can replicate complex structures of a native ECM and allow structural properties to emerge. Previous research on photopatterning showed the potential of tuning biophysical and biochemical cues in patterned materials (25). To study the effect of stiffness and degradation on 3D cell behavior, we use the combination of two different types of crosslinking. The first type of crosslinking is covalent Diels-Alder click chemistry, which offers an efficient and versatile reaction for hydrogel formation (11)(13). The second type of crosslinking, UV-mediated thiol-ene peptide binding, offers tunable degradability by the matrix metalloprotease enzymes secreted by encapsulated cells (16). Despite the numerous research performed on single-phase materials, fewer investigations are looking at cell response in multicomponent matrices such as patterned materials.

Dual crosslinked, patterned hydrogels previously described have shown an effect on cells attached to 2D substrates, such as in cell alignment (26), protein expression and differentiation (27)(26). Previous research in 3D cell encapsulation showed that patterns in biochemical cues can influence cell migration (28) and localized growth (29), whereas patterns in biophysical cues can influence cell interactions (30). Research performed on patterning multiple mechanical or biochemical characteristics has shown promising results on guiding cell behavior (31). Our research contributes on evaluating the cell response in patterned hydrogels with spatially discrete patterns in degradation and stiffness. Furthermore, the evaluation of the cell response in patterned materials has been limited to the independent evaluation of each phase; no method has been proposed to quantitatively assess patterned cell response in a multicomponent matrix. To achieve this, an image-based analysis tool is required.

Here we present an alginate-based hydrogels with anisotropic stiffness-degradation spatial patterns and compatible with 3D cell encapsulation. The hydrogels exhibit emerging patterns in stiffness and degradability over time, taking advantage of dual covalent Diels-Alder click crosslinking and UV-mediated peptide crosslinking. Further, we develop a novel quantitative, image-based analysis tool to evaluate the emerging anisotropic cell behavior in 3D and over time. We characterize cell morphology and proliferation in photopatterned materials and compare the results with equivalent single-phase materials. Such patterned materials allowing the emergence of 3D anisotropic cell response, together with the image-based analysis method, are valuable tools to understand cell-matrix interactions in multicomponent materials.

## 2 Materials and Methods

### 2.1 Alginate modification

To form the click-crosslinking, norbornene and tetrazine must be added in the alginate backbone. The alginate used was low molecular weight, high guluronic acid sodium alginate (MW 75kDa Pronova UP VLVG; NovaMatrix). The coupling of norbornene (N, TCI Chemicals, #N0907) and tetrazine (T, conju-probe, #CP-6021) to the alginate molecule was performed as previously described (27). Alginate modification with norbornene was performed with a theoretical degree of substitution (DS_theo_) of DS_theo_ 200 for norbornene. Tetrazine modification was performed with a DS_theo_ 50 for tetrazine. To determine the reaction efficiency and the actual DS (DS_actual_) required to ensure appropriate norbornene to tetrazine (N:T) ratios for crosslinking, NMR measurements were performed, using a 1.5% w/v alginate solution in deuterium oxide (64 scans; Agilent 400 MHz Premium COMPACT equipped with Agilent OneNMR Probe) and analyzed using MestrNova Software (14.6) (Supplementary Figure S1; Supplementary Table S1).

### 2.2 Mouse Embryonic Fibroblast (MEF) cell culture

Mouse embryonic fibroblasts (SCRC-1040; ATCC) were cultured in Dulbecco’s Modified Eagle’s Medium (Sigma, #D5546) supplemented with 3.5 g/l glucose (VWR, # 0188), 15% v/v fetal bovine serum (Biochrom, #S0615), and 1% penicillin/streptomycin (Gibco, #15140-122). Cells were maintained in a 5% CO_2_ environment at 37°C and passaged every 3–5 days. For 3D encapsulation, cells were used at passage 16.

### 2.3 Hydrogel formation

The hydrogel formation was performed based on previously established protocols (27) with modifications in N:T ratios and alginate concentration, as described below.

#### 2.3.1 Non-degradable matrix: Click-crosslinked hydrogels

The precursors for the hydrogel were dissolved in phosphate-buffered saline (PBS, without Ca^2+^, Mg^2+^ and phenol red; Biozym) and distributed into 2 tubes. The first tube contained norbornene-modified alginate (N-alg); MMP-sensitive (MMPsens) peptide (GCRD-VPMS ↓ MRGG-DRCG, 98% purity; WatsonBio) at a final concentration of 10 mg/ml of hydrogel, thiolated RGD-peptide (CGGGGRGDSP; Peptide2.0) at a concentration of 5 molecules of RGD per alginate chain (DS 5), and the cell suspension at final concentration of 5×10^6^ cells/mL of hydrogel. The second tube contained tetrazine-modified alginate (T-alg) and the photoinitiator (Irgacure 2959; Sigma-Aldrich, #410896) at a final concentration of 3 mg/mL of hydrogel. The total final concentration of alginate was 2% w/v at an N:T ratio of 1.5.

The two solutions were mixed by pipetting and cast onto a bottom glass plate, with the casting area being restricted on three sides by glass spacers, and immediately covered with a glass slide previously treated with SigmaCoat (≥99.5%; Sigma-Aldrich, #SL2) to prevent adhesion. The gel height was constrained to 2 mm by the thickness of the glass spacers. Spontaneous click-crosslinking for 50 min at room temperature (RT) and in the dark allowed the N:T covalent bonds to form. Despite MMPsens and the photoinitiator being present, these were not activated due to the lack of UV exposure. Nevertheless, the MMPsens and photoinitiator need to be present to allow for patterned materials (see section 2.3.3).

In order to ensure a homogeneous binding of the RGD-peptide, crosslinked gels were exposed to 2 min UV light (365 nm) at 10 mW/cm^2^ (Omnicure S2000) in a custom-built exposure chamber. The cylindrical hydrogels were punched from the cast gel sheet using 5 mm biopsy punches (Integra Miltex) and placed in growth media at 37°C and 5% CO_2_.

#### 2.3.2 Degradable matrix: MMPsens peptide crosslinked hydrogels

The production of degradable materials followed the same procedure as described in section 2.3.1, with an additional step for the MMPsens peptide crosslinking. After casting the hydrogel solution between the glass plates, the material was exposed to UV light at 10 mW/cm^2^ for 10 min to initiate the coupling of the degradable MMPsens peptide to the norbornene-modified alginate via thiol-ene crosslinking. After the UV exposure, the materials were placed for an additional 50 min at RT in the dark to allow for the N:T covalent bonds to be formed. To ensure a homogenous binding of RGD, the hydrogels were exposed again to UV for 2 min. Hydrogels were punched out and incubated in growth media at 37°C and 5% CO_2_.

As negative control materials, hydrogels were fabricated with peptide crosslinkers not susceptible to degradation, MMP-scramble (VpMSmRGG). In this case, the peptide contained the same sequence as the degradable isoform but with some amino acids in the D-form (indicated in lower case letters), rendering them unrecognizable to matrix metalloprotease enzymes.

#### 2.3.3 Patterned: Dual crosslinked hydrogels

The creation of patterned materials followed the same procedure as described in section 2.3.2, with the addition of a photomask placed on top of the cover glass during the UV mediated thiol-ene coupling of the MMPsens peptide. The photomask had a pattern of straight lines with 500 μm thickness (UV light blocking sections, non-degradable matrix equivalent to 2.3.1) placed 250 μm apart (UV light permitting sections, degradable matrix equivalent to 2.3.2).

### 2.4 Mechanical characterization

Mechanical characterization was performed on day 1 and day 14. All mechanical characterization was performed with cell-loaded materials to quantify the enzymatic degradation of the hydrogels in stiff and degradable (Stiff-Deg) materials. This was also true for soft and non-degradable (Soft-NoDeg) materials to keep comparable conditions. The material degradation was evaluated via three different methods: unconfined compression testing for measuring bulk elastic modulus of single-phase materials, rheology to quantify loss and storage modulus of single-phase materials and microindentation to estimate the surface elastic modulus of single-phase and patterned materials.

### 2.5 Unconfined compression testing

Single-phase materials were subjected to uniaxial unconfined compression testing (BOSE Test Bench LM1 system) with a 250 g load cell (Model 31 Low, Honeywell) at 0.016 mm/s without preload. The elastic modulus *E* was calculated as the slope of the linear region of the generated stress vs. strain curve, in the 2-10% strain range, using a MATLAB (R2019b) script (n = 6). The required MATLAB inputs of hydrogel height and diameter were determined by lowering down the BOSE system top plate until contact with the gel surface was established and by using calipers, respectively.

### 2.6 Rheology

Storage and loss modulus of single-phase hydrogels were determined with a rheometer (Anton Paar MCR301) via frequency sweeps with a parallel plate geometry of 8 mm (PP08, Anton Paar). The frequency sweep was performed from 0.01 to 10 Hz and at 0.1% shear strain at RT (n = 6). Once contact with the gel surface was established, a pre-compression of 10% of the height of the hydrogel was applied prior to the measurement. No additional hydration was needed as the experiment lasted less than 10 min. To obtain the elastic modulus, first the shear modulus (G) was derived from the storage (G’) and loss (G”) modulus using Rubber’s elasticity theory (Eq. 1).

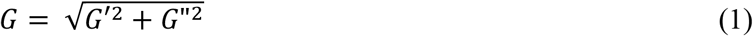

The elastic modulus (E) was calculated using the values of the shear modulus obtained from Eq.1 (32) and the approximation of Poisson’s ratio (υ) equal to 0.5 (33) (Eq. 2).

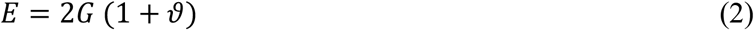

The mesh size (ξ) was approximated by Eq. 3, proposed for alginate hydrogels, in which the storage modulus G’ in low frequencies (0.1-1Hz) was used (34), with *N*_*aν*_ being avogadro’s number (6.022 10^23^ 1/mol), *R* being the ideal gas constant (8,314 m^3^Pa/K° mol) and *T* being the room temperature (293°K).

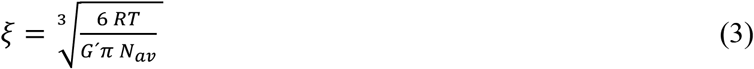

### 2.7 Microindentation

#### 2.7.1 Depth-Sensing Indentation/Air-Indent Method

Depth-sensing microindentation measurements were done using a Triboindenter TI-950 (Hysitron-Bruker, MN, USA) equipped with an XZ-500 extended displacement stage, allowing a vertical displacement of up to 500 μm (35). After the first contact to detect the surface, the tip was retracted for ~300 μm. Next, the measurements were conducted using the “air-indent” mode, allowing a reliable indentation curve without any additional sample pre-contact. The measurements were done using a cono-spherical tip of 50 μm radius and in automated mode to map an area of 6×6 matrix, indentation spacing of 300 μm in single-phase materials and 18×11 matrix with an indentation spacing of 150 μm in patterned materials. The measurements were done in displacement control mode, using a displacement function of 250 μm retraction and 300 μm approach, with a strain rate of ~30 μm/s.

#### 2.7.2 Analysis of load-displacement curves

To meet the Hertzian contact model requirement, the first 30 μm of contact depth after initial contact, in which the tip geometry stays spherical, was used for curve fitting and calculation of the indentation elastic modulus (Eq. 4). This model was chosen as it describes the contact mechanics of 3D solids and correlates the elastic modulus (*E*) with the contact surface radius (*R*, 50 μm), load (*y*) and contact depth (*x*)

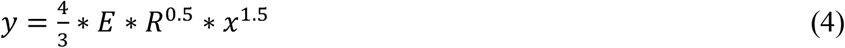

Considering the high number of indents, the analysis of the load-displacement curves was automated by a custom-made Python3 script. The depth of the gel and the load of the indenter (both ordered by time) are the main data vectors used for the analysis. This automation is divided into four main parts: (1) identifying the point of interest (POI), (2) extracting the curve segment, (3) fitting the Hertzian model on the extracted segment and (4) obtaining the indentation *E* value per indentation point, collected in a matrix and depicted in a heat map. Further information can be found in Supplementary Figure S2.

### 2.8 Cell viability by Live/Dead staining

Cell viability was assessed after 1 and 14 days using Live/Dead staining. The hydrogels were taken out from the incubation media and washed with PBS. Then the cells were stained with a solution of 4 mM calcein AM (TRC, #C125400) and 4 mM ethidium homodimer-1 (Thermo Fisher, #L3224) dissolved in PBS to identify live and dead cells, respectively. The staining solution volume was 400 μl per hydrogel, stained for 12 min in a cell culture incubator at 37°C, 5% CO_2_ in the darkness. A final washing step was performed with 400 μl of PBS per hydrogel at RT for 5min and protected from light.

Imaging was performed on a confocal microscope (Leica SP5, Germany). Quantification of cell number and viability at each time point was performed using ImageJ software (ImageJ 1.53s) (36). Three independent positions per gel were acquired at the gel center at 25x magnification, from 2 independent samples, resulting in n=6 fields of view containing multiple single cells (n>100). To assess cell proliferation (cell number per unit volume), differential swelling of soft and stiff hydrogels was taken into account, as explained in Supplementary Information S3.

### 2.9 Cell morphology by DAPI/Phalloidin staining

To evaluate cell morphology, DAPI/Phalloidin staining was performed after 1 and 14 days, visualizing nuclei and actin, respectively. All steps were performed under orbital shaking, in a 24 well plate and using a volume of 400 uL per gel. Encapsulated cells were fixed in 4% paraformaldehyde solution (Sigma Aldrich, Sigma, #158127) for 45 min at RT, then permeabilized with 0.3% Triton X-100 (Sigma Aldrich, #11488696) for 15 min, washed twice with 3% bovine serum albumin (BSA, Sigma, #A2153) in PBS for 5 min and stained in the dark with 4, 6-diamidino-2-phenylindole (DAPI; Sigma, #MBD0015) and TRITC-conjugated Phalloidin (Cell Signaling, #8878S) for 3h. A final wash was performed with 3% BSA in PBS for 5 min at RT.

Three independent positions per gel were acquired at the gel center using a confocal microscope (Leica SP5, Germany). For a general quantification of cell morphology, 25x magnification was used and n=6 fields of view (3 different images from 2 independent samples) were taken, containing multiple single cells (n>50). In addition, ten single cell images per gel (5 cells from 2 different hydrogels) were analyzed for quantification of filopodia number and length. Images were obtained from the center of the gel using 64x magnification.

### 2.10 Image-based analysis tool to study anisotropic multicomponent materials

A custom-made image-based analysis tool in the form of a macro written in ImageJ (ImageJ 1.53s) (36) has been created to analyze cellular readouts obtained from Z-stack projections from anisotropic patterned materials. The macro offers the possibility to freely divide an image into rectangular units, which leads to a heat map in which the results are later depicted. The background is separated from the cells via a threshold. To compensate for pixel noise from the raw data, a denoise function (median filter) is built in, which can be used with different strengths depending on the image. In this way, a binary mask is created, which is used for most of the calculations. For details on the binning size optimization, refer to Supplementary Figure S4.

Three readouts are calculated for every tile within the heat map: Cell Projected Area, Cell Circularity and Cell Number. Cell Projected Area is calculated for each cell as number of pixels and converted into μm^2^ or mm^2^. Cell Circularity is calculated for each cell as 4π*area/perimeter^2, where 1 indicates a perfect circle and values towards 0 indicate elongated cells. Cell Number is calculated as number of DAPI nuclei within each tile. Every cell in a tile will be individually calculated and the mean of all cells in a tile is used. Cells touching the tile border are excluded. For further details, refer to Supplementary Figure S5.

### 2.11 Statistical analysis

Results are depicted as bar graphs with mean and standard deviation, or box plots with median, 1^st^ and 3^rd^ quartile, using OriginLab (Pro 2022b). Comparison of hydrogel mechanical properties were performed using Student t-test (p < 0.05). Comparison of cellular read-outs were performed using Student t-test (p < 0.05) for normally distributed data and Wilcoxon Signed Rank test (p < 0.05) for not normally distributed data.

## 2 Results

### 2.10 Mechanical characterization

Single-phase Stiff-Deg and Soft-NoDeg materials were characterized for their bulk elastic and viscoelastic properties at day 1 and day 14, as well as changes over time, using rheology and unconfined compression testing. The storage modulus (G’) of Stiff-Deg is higher than Soft-NoDeg materials with average values of 3353 ± 36 Pa and 530 ± 10 Pa, respectively, at day 1 (Fig. 1A) and 1848 ± 41 kPa and 776 ± 26 kPa at day 14 (Fig. 1B). The values of G’ showed a decrease at day 14 (Fig. 1B) compared to day 1 (Fig. 1A) for Stiff-Deg materials, whereas G” modulus presented a similar behavior at day 1 and day 14 for both materials.

**Figure 1:**
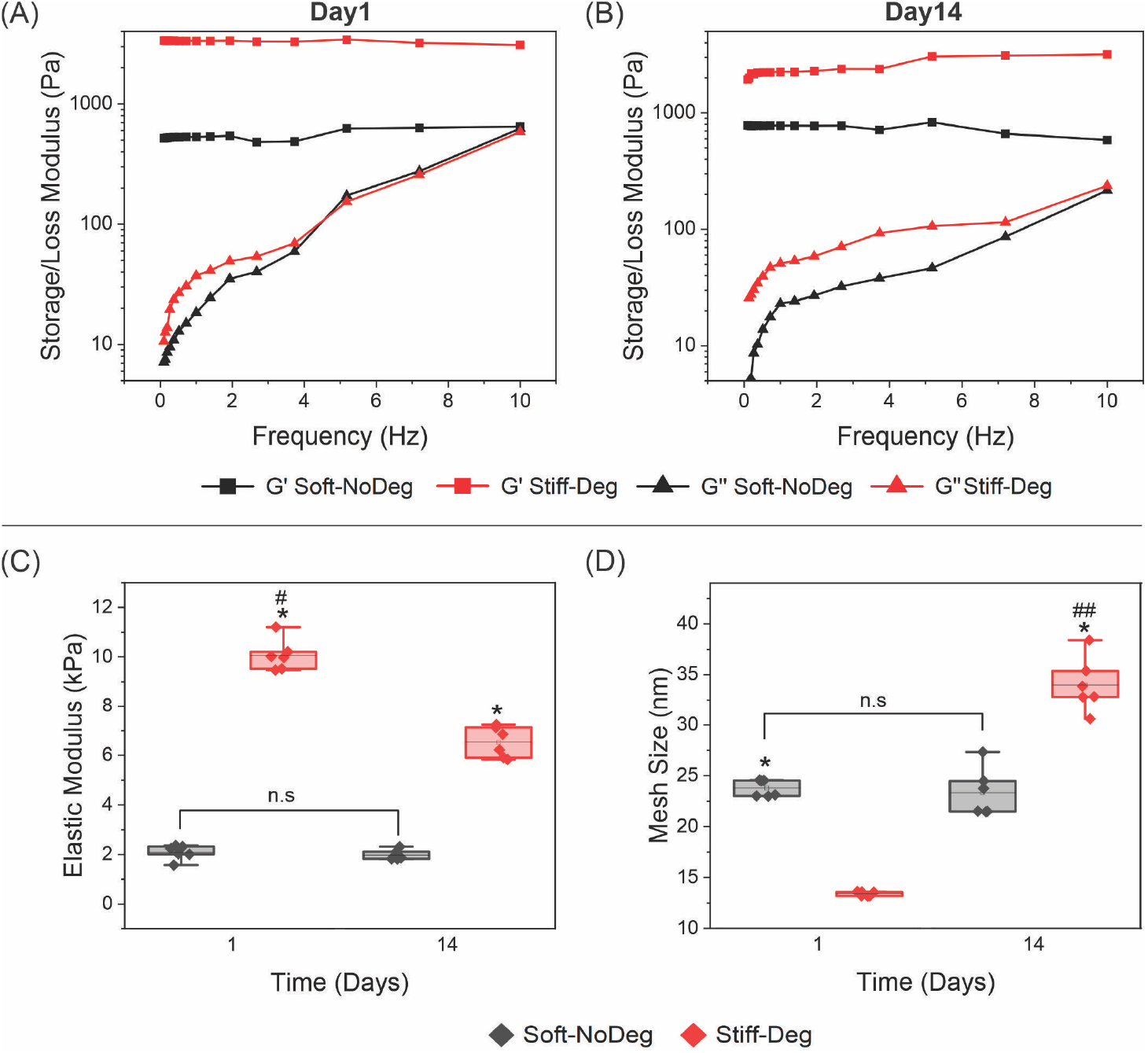
Mechanical characterization of single-phase materials: Soft-NoDeg (black) and Stiff-Deg (red). (A) Day 1 and (B) day 14 of storage (G’, □) and loss (G”, Δ) modulus in Pa obtained by rheology, n=6 gels. (C) Elastic modulus determined by unconfined compression testing in kPa, n=6 gels. (D) Mesh size estimated from the storage modulus in nm, n=6 gels. Statistical significance with Student t-test for differences between groups is indicated with * and differences between time points with # (*/# = p<0.05, **/## = p<0.01).

Bulk elastic modulus was characterized by unconfined compression testing (Fig. 1C). At day 1, there is a significant difference between the Soft-NoDeg (2 ± 0.3 kPa) and Stiff-Deg (10 ± 0.6 kPa) materials. At day 14, there is a significant decrease of elastic modulus in Stiff-Deg materials (6 ± 0.6 kPa) with respect to day 1. The Soft-NoDeg materials showed a constant elastic modulus at day 14 (2 ± 0.2 kPa).

The dynamic behavior of degradable materials is also evident in the change of the mesh size (Fig. 1D). The mesh size increases significantly in degradable materials from 13.0 ± 0.1 nm on day 1 to 34 ± 3 nm on day 14. In contrast, Soft-NoDeg materials maintain the mesh size over 14 days, as the values of day 1 (24 ± 0.3nm) and day 14 (26 ± 2 nm) are not significantly different.

To characterize the anisotropic mechanical properties of patterned hydrogels we used the method of microindentation. Patterned materials show a clear difference in the elastic modulus between the 2 phases, on day 1 (Fig. 2A) and day 14 (Fig. 2D). The corresponding single-phase materials showed similar values of elastic modulus. The surface elastic modulus of Soft-NoDeg materials was comparable between day 1 (Fig. 1B) and day 14 (Fig. 1E) and the elastic modulus of the Stiff-Deg materials decreased visibly between day 1 (Fig. 1C) and day 14 (Fig. 1F).

**Figure 2:**
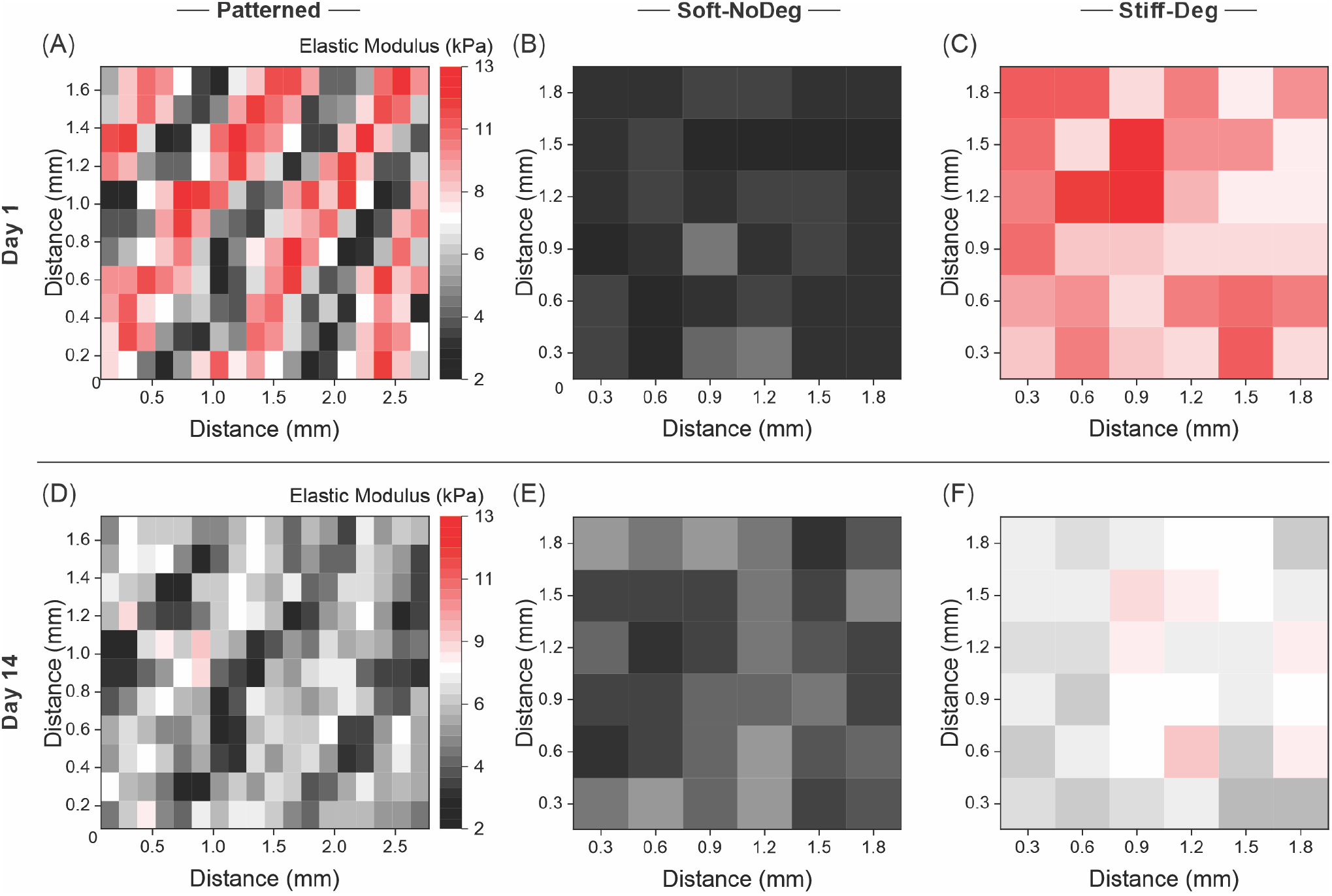
Microindentation of single-phase and patterned materials. (A, D) Patterned materials, (B, E) Soft-NoDeg single-phase materials and (C, F) Stiff-Deg single-phase materials, on day 1 and day 14, respectively. Each matrix is the visual representation of the indentation elastic modulus (kPa) at the material surface. Single-phase materials (6×6 matrix, indentation spacing of 300 μm), patterned materials (18×11 matrix, indentation spacing of 150 μm).

### 3.2 Cell viability and proliferation in 3D single-phase and patterned materials

Mouse embryonic fibroblasts were encapsulated in 3D single-phase and patterned hydrogels. Cell viability was evaluated at day 1 and day 14 by staining live cells with calcein (green) and dead cells with ethidium homodimer-1 (red).

Single-phase materials showed high viability (Fig. 3A), as the fraction of viable cells remained above 90% for all materials and time points. The cell number corrected to the swelling factor (Fig. 3B) shows that the cell proliferation was higher in Stiff-Deg materials compared to Soft-NoDeg, with significantly higher cell number at day 14 compared to day 1 and compared to the Soft-NoDeg counterpart at day 14. In contrast, no significant differences over time were seen in the cell number for Soft-NoDeg materials.

**Figure 3:**
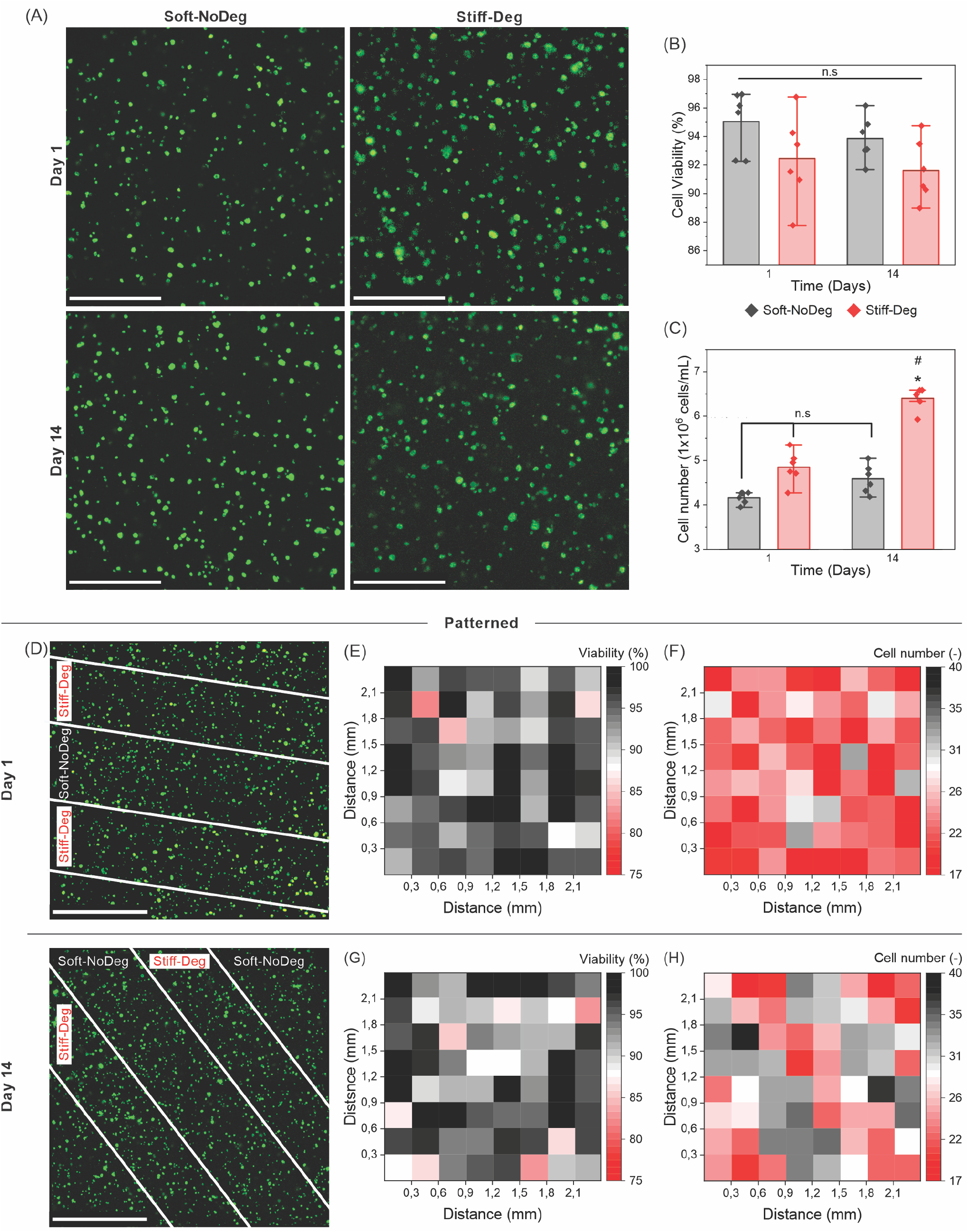
Viability and proliferation of encapsulated cells in single-phase and patterned materials on day 1 and day 14. (A) Live/Dead staining of Soft-NoDeg and Stiff-Deg single-phase materials at day 1 and day 14, 25x magnification, 250 μm z-stack, and corresponding (B) cell viability in % (viable cells/total cells) and (C) cell number (cells per mL of hydrogel). (D) Live/Dead staining of patterned materials at day 1 and day 14, with 2 × 2 tile merging of 10x magnification, 250 μm z-stack. The macro function “cell number” was used to quantify and plot the heat maps corresponding to cell viability in patterned materials at (E) day 1 and (G) day 14, as well as total cell number at (F) day 1 and (H) day 14. The bars in B and C represent the mean and standard deviation of n = 6 fields of view containing multiple single cells (n>100). Statistical significance with Student t-test for differences between groups is indicated with * and differences between time points with # (*/# = p<0.05, **/## = p<0.01). Scale bar: 500 μm (A), 1 mm (D).

The macro function “cell number” allowed the quantification and visualization of cell viability and proliferation in patterned materials. Comparable to single-phase materials, patterned materials also showed high viability in both phases and over time (Fig. 3D). No visible patterns or changes were shown in viability, neither at day 1 (Fig. 3E) or day 14 (Fig. 3G).

Encapsulated cell number showed an initial homogeneous distribution of cells, as on day 1 there are no visible patterns (Fig. 3F). However, patterns in cell proliferation are evident at day 14, which show higher cell number in the Stiff-Deg areas compared to the Soft-NoDeg zones (Fig. 3H).

### 3.3 Cell morphology in 3D single-phase materials

Staining of the nuclei (DAPI, cyan) and the actin cytoskeleton (phalloidin, green) in single-phase materials was used to analyze the effect of material properties on cell morphology.

On day 1, cells in Soft-NoDeg materials displayed significantly greater projected area compared to cells in Stiff-Deg materials (Fig. 4B). 14 days after encapsulation, when the Stiff-Deg materials degraded and consequently softened, the cell projected area increased significantly compared to the initial time point and also in comparison with the Soft-NoDeg materials at day 14.

**Figure 4:**
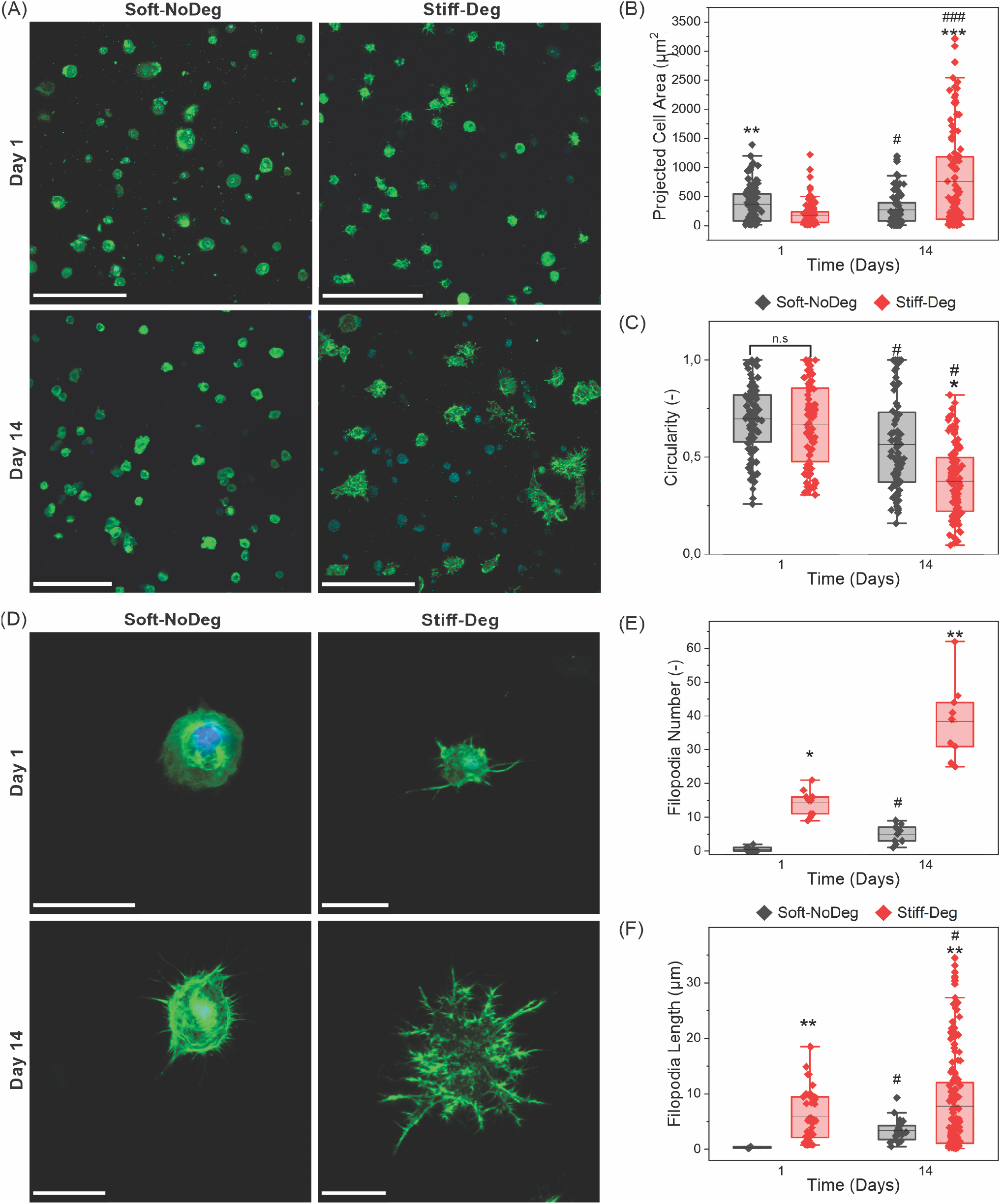
Morphology of encapsulated cells in single-phase materials at day 1 and day 14. (A) Phalloidin (green)/ DAPI (cyan) staining of multiple cell images with 25x magnification, 250 μm z-stack to determine (B) projected cell area in μm^2^ and (C) circularity (−). (D) Higher 40x magnification of single cell z-stack to determine (E) filopodia number (−) and (F) filopodia length in μm. Boxes represent the median and 1^st^ and 3^rd^ quartile of (B, C) multiple cells (>50cells) in n=6 fields of view or (E, F) n=10 cells. Statistical significance with Wilcoxon Signed Rank test for differences between groups is indicated with * and differences between time points with # (*/# = p<0.05, **/## = p<0.01). Scale bar: 200 μm (A), 25 μm (D).

Differences in cell circularity at day 14 are significant between the 2 materials (Fig. 4C). The cells in Stiff-Deg materials show significantly lower circularity compared to the initial time point and to cells in Soft-NoDeg hydrogels at day 14.

In Figure 4D, single cell images are shown, depicting detailed cell morphology and filopodia. On day 1, early filopodia formation can be seen in Stiff-Deg materials, whereas no filopodia were formed in Soft-NoDeg hydrogels. After 14 days, the filopodia number and length increased significantly in Stiff-Deg compared to the initial time point and to Soft-NoDeg at day 14. In Soft-NoDeg materials, filopodia number and length increased after 14 days of encapsulation, yet they remained lower compared to Stiff-Deg materials.

### 3.4 Cell response in 3D patterned materials

The photopatterning of single-phase materials created anisotropic hydrogels with spatially distinct degradation and stiffness characteristics.

Figure 5 shows the effect of patterned materials on the morphology of MEFs (Fig. 5A-F), the evaluation and heat map representation using the novel image-based analysis tool (Fig. 5G-J) and the quantification of the individual material phases (Fig. 5K-N). On day 1 (Fig. 5A, C, E), there are patterns in projected cell area (Figure 5G) as the Soft-NoDeg phase shows cells with significantly larger projected cell area compared to Stiff-Deg (Fig. 5K). Initially, no significant patterns in circularity are visible (Fig. 5H, L) as most of the cells present a round morphology. At day 14 after encapsulation (Fig. 5B, D, F), there is a significant increase of the projected cell area in the Stiff-Deg (Fig. 5K) and even stronger significant decrease in cell circularity (Fig. 5L). This is visualized in the heat maps with emerging spatial patterns in cell circularity at day 14 compared to day 1 (Fig. 5J, H) and less visible, even reverted patterns in projected cell area (Fig. 5I, G).

**Figure 5:**
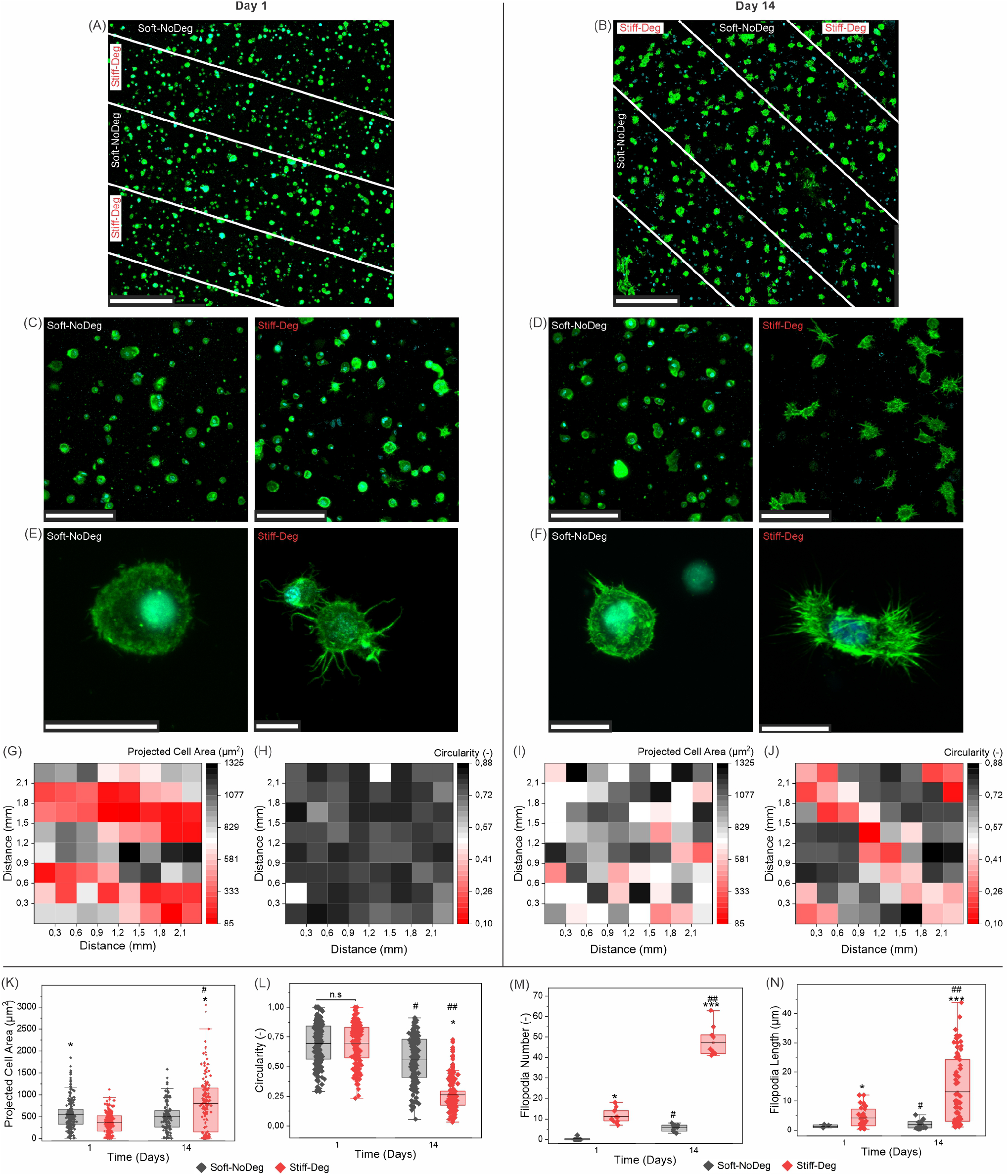
Morphology of encapsulated cells in patterned materials at day 1 and day 14. Phalloidin (green)/ DAPI (cyan) staining overview images at (A) day 1 and (B) day 14, with indicated pattern areas, 2×2 tile image, 10x magnification, 250 μm z-stack. Zoom-in on the individual regions of the pattern at (C) day 1 and (D) day 14, 25x magnification, 250 μm z-stack. Single cell images z-stack at (E) day 1 and (F) day 14, 40x magnification. Heat map representation of (G, I) the mean projected cell area in μm^2^ and (H, J) circularity (−) in the overview images, at day 1 (G, H) and day 14 (I, J). Box plots quantifying (K) projected cell area, (L) circularity, (M) filopodia number (−) and (N) filopodia length in μm, at day 1 and day 14, showing the median and 1^st^ and 3^rd^ quartile of n=6 fields of view containing multiple single cells (n>50, for K and L) or n=10 cells (for M and N) in patterned materials. Statistical significance with Wilcoxon Signed Rank test for differences between groups is indicated with * and differences between time points with # (*/# = p<0.05, **/## = p<0.01). Scale bar: (A, B) 500 μm, (C, D) 200 μm, (E, F) 25 μm.

Regarding cell morphology, single cell images at day 1 (Fig 5E) showed that filopodia are mainly formed in Stiff-Deg phase, with significantly greater number (Fig. 5M) and length (Fig. 5N) of the filopodia. This trend is amplified at day 14 (Fig 5F), with significantly increased filopodia number and length compared to day 1 and compared to cells in Soft-NoDeg phase.

## Discussion

The presented 3D hydrogels with stiffness-degradation spatial patterns allow cell encapsulation with high cell viability and anisotropic cell response. The hydrogel casting procedure offers the possibility of photopatterning, combining the properties of two single-phase materials in one single, multicomponent matrix, which allows emerging patterns in cell behavior in 3D. Evaluation of cell behavior in multicomponent materials is crucial in order to understand how these platforms guide cell response. In our case, we choose patterns in stiffness-degradation and evaluate anisotropic fibroblast cell morphology, as an example of the application of an image-based quantification method.

All methods used for mechanical characterization led to consistent and comparable results of mechanical properties and changes over time caused by degradation. First, the methods show a decrease over time of the elastic modulus of Stiff-Deg materials compared Stiff-NoDeg materials. Second, the bulk elastic modulus of the single-phase materials is comparable to the surface elastic modulus of single-phase materials, and importantly, also consistent with the mechanical properties of the respective phases of patterned multicomponent materials.

The decrease in the elastic modulus of the degradable material can be attributed to the degradation of the MMPsens peptide bonds due to the action of the enzymes secreted by the cells. A consequence of this degradation can be shown in the significant increase of the mesh size over time. There is no significant change in the mesh size of Soft-NoDeg materials, as the covalent bonds of these hydrogels are non-degradable.

Our results showed that the projected cell area of 3D encapsulated cells is dependent on the matrix stiffness. At day 1, the significantly lower elastic modulus of Soft-NoDeg vs. Stiff-Deg results in significantly higher projected cell area in both single-phase and patterned materials. However, at day 14, when the elastic modulus of Stiff-Deg significantly drops compared to day 1, the projected cell area significantly increases and cell circularity decreases as degradation promotes cell spreading. These results are supported by previous results related to 3D fibroblast encapsulation (37) and in contrast to cell behavior on 2D surfaces with patterns in stiffness (38), as expected.

Matrix remodeling and dynamic environments are crucial to stimulate cell response (39). Degradation is essential for the formation of protrusions and we observe that Stiff-Deg materials promote longer and higher filopodia number compared to Soft-NoDeg materials. The control hydrogels formed with a non-degradable version of the peptide (MMP-scramble), showed that cells do not form filopodia in non-degradable materials (Supplementary Figure 6). These results are supported by previous findings on the effect of matrix deformation energy in the actin cytoskeleton of the cell, which has been proven to have a greater effect compared to the intrinsic matrix stiffness (40). Such findings highlight the importance of matrix degradability in enabling cell protrusions to invade into the surrounding environment, as they regulate more advanced cell processes like migration, motility, communication and differentiation (41).

One important feature of this work is the combination of Stiff-Deg and Soft-NoDeg phases in one single, multicomponent matrix. Differences in cell response observed in single-phase materials are recapitulated in patterned stiffness-degradation materials and, importantly, anisotropic cell behavior emerges with time as the Stiff-Deg component degrades. This sets the basis for future work looking at sharper material interfaces, or in contrast, gradients of stiffness-degradability by manipulating the photomask. Such multicomponent materials open opportunities to investigate anisotropic 3D cell migration, proliferation or differentiation across a cell-relevant stiffness-degradability range.

To evaluate anisotropic 3D cell response in patterned materials, we have developed a new image-based analysis tool and visual presentation of spatial anisotropies of material and cellular characteristics using heat maps. Various research groups have evaluated patterned materials as independent phases, not as a single, multicomponent matrix. The developed image-based method and the heat map representation of cell number and morphology (projected cell area and circularity) showed to be a valid tool to characterize and quantify anisotropic 3D cell behavior in patterned materials, as it consistently represented the anisotropic cell behavior in each phase compared to corresponding single-phase controls. This image-based analysis could be extended to other image-based cellular read-outs.

Despite the great advantage of our novel image-based analysis tool, there are some limitations. As input for this analysis tool, images covering the entire gel or stitched multi-tiles images are required. However, for certain features such as filopodia formation, high magnification images are necessary. Multi-tiles high magnification imaging covering the entire gel currently requires long acquisition times, which would lead to dehydration of the hydrogel.

Our research demonstrates a relevant approach to investigate emerging anisotropic 3D cell behavior in stiffness-degradation patterned materials. The developed image-based analysis method provides the basis for visualizing and quantifying 3D anisotropic cell behavior with regard to cell number, cell projected area and circularity. This anisotropic 3D cell response was confirmed with high resolution quantification of filopodia number and length. Such stiffness-degradation patterned hydrogels allowing the emergence of 3D anisotropic cell response, together with the image-based analysis method for visualization and quantification of cellular read-outs, are valuable tools to understand cell-matrix interactions in multicomponent materials.

## 3 Conflict of Interest

*The authors declare that the research was conducted in the absence of any commercial or financial relationships that could be considered as a potential conflict of interest*.

## 4 Data availability

All raw and processed data, and the MATLAB and Python scripts are available in a publicly accessible repository of the Max Planck Society https://doi.org/10.17617/3.NEHZN1.

## 5 Author Contributions

A Cipitria conceived the idea. CA Garrido and DS Garske performed the experiments. S Amini supported the microindentation experiments. S Real developed the algorithm for analysis of the microindentation data. CA Garrido quantified and analyzed the data. M Thiele developed the image-based analysis macro. K Schmidt-Bleek and GN Duda evaluated the methods and results. CA Garrido and A Cipitria drafted the manuscript. All authors discussed the results and contributed to the final manuscript.

## 6 Funding

This work was funded by the Deutsche Forschungsgemeinschaft (DFG) CRC 1444 grant. A. C. also thanks the funding from the DFG Emmy Noether grant (CI 203/2-1), IKERBASQUE Basque Foundation for Science and from the Spanish Ministry of Science and Innovation (PID2021-123013OB-I00).

## 7 Acknowledgments

The authors acknowledge the support from all group members of Cipitria, Schmidt-Bleek and Duda’s laboratories.

## Notes

### Competing Interest Statement

The authors have declared no competing interest.

