## Supplementary information for "Hydrogels with stiffness-degradation spatial patterns control anisotropic 3D cell response"


- Supplementary Figure S1
- Supplementary Table S1
- Supplementary Figure S2
- Supplementary Information S3
    - Supplementary Figure S3
    - Supplementary Table S3
- Supplementary Figure S4
- Supplementary Figure S5
- Supplementary Figure S6

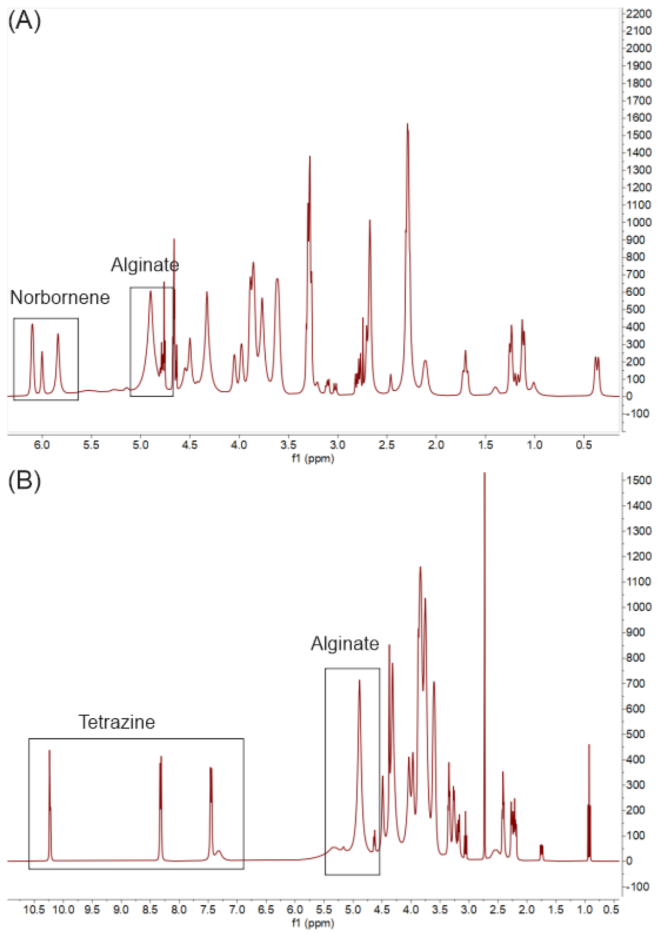


**Supplementary Figure S1:** NMR spectra of modified VLVG alginate. (A) NMR of norbornene modified alginate with the 3 characteristic peaks of norbornene between 6.2-5.8 ppm. (B) NMR of tetrazine modified alginate with the 3 characteristic peaks of tetrazine at 10.2, 8.2 and 7.4 ppm.

| | Norbornene | Tetrazine |
|---|---|---|
| **$DS_{theo}$** | 200 | 50 |
| **$DS_{actual}$** | 43.7 | 40.8 |
| **Reaction efficiency (%)** | 21.8 | 81.6 |

**Supplementary Table S1:** Norbornene (N-Alg) and tetrazine (T-Alg) modified alginate with a theoretical degree of substitution ($DS_{theo}$), actual DS ($DS_{actual}$) and reaction efficiency (%).

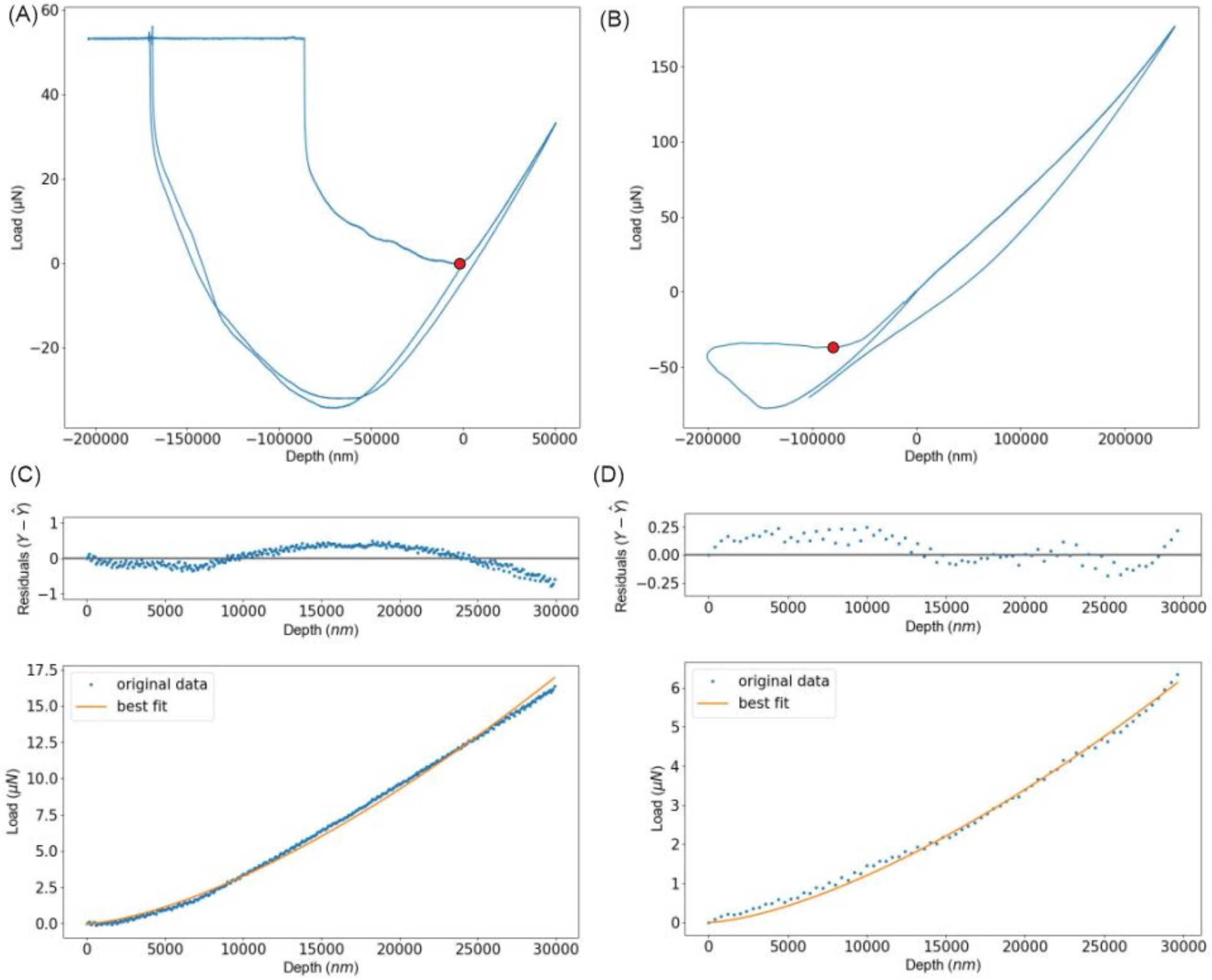


**Supplementary Figure S2:** Determination of the elastic modulus of patterned hydrogels via surface microindentation using a custom-made Python3 script for the quantification workflow. The raw data obtained from the microindentation is plotted as depth vs. load. The first step is to determine the origin point (point of interest, POI), shown as a red dot for (A) soft and (B) stiff materials. This is achieved by 2 different methods: (i) a fast method that detects the first point, determined as the minimum where a decrease is followed by an increase of values. An alternative method (ii) uses a minimum slope value of a linear regression created by a group of points, which allows selecting a specific slope value. The second step is to extract from the curves in (A) and (B), the segment corresponding to the first 30 μm of indentation (30000 nm on x axis after the origin, shown in the blue lines of (C) and (D)) and fit to the Hertzian approximation (orange lines of (C) and (D)). In order to verify the approximation, the values of the residuals are plotted, as a comparison of y value real ($Y$) and the y value of the regression ($\hat{Y}$). Finally, the output file is a matrix with the elastic modulus values obtained from the regression curves.

**Supplementary Information S3:** Calculation of the cell number per unit volume taking into account the differential swelling of hydrogels with different mechanical and degradation properties

The hydrogels swell over time (example in Fig. S3.A and B), influencing the cell count in the field of view of the microscope (see Fig. S3.C).

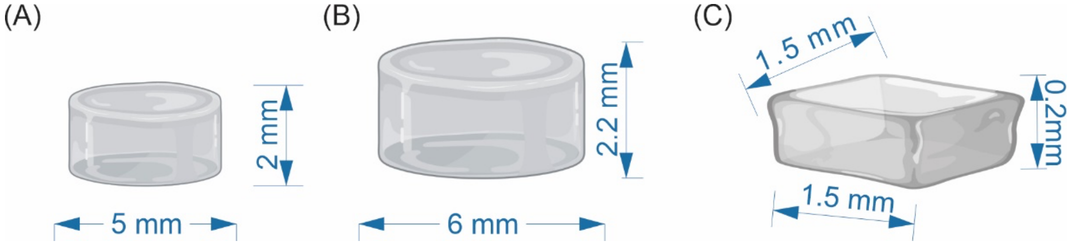


**Supplementary Figure S3:** Examples of dimensions of hydrogels and the microscope field of view. (A) Hydrogel dimensions at day 1. (B) Hydrogel dimensions at day 14. (C) Field of view used for calculating cell number per unit volume with 10x magnification.

The cell number per unit volume ($C_V$) is calculated by counting the cells in the field of view and dividing by the volume of the z-stack (Fig. S3.C). At day 1, no corrections are required. However, at day 14, the cell number per unit volume needs to be corrected ($C_V^*$) due to differential swelling, particularly for highly swelling materials. We do so multiplying the cell number per unit volume ($C_V$) by the volumetric swelling, where $V_{day\ 14}$ is the overall gel volume at day 14 and $V_{day\ 1}$ is the overall gel volume at day 1.

$$C_V^* = C_V \frac{V_{day\ 14}}{V_{day\ 1}} \qquad \text{(S Eq. 1)}$$

The overall gel volume is measured using calipers. The average volumetric swelling of Soft-noDeg and Stiff-Deg materials are shown in Supplementary Table S3.

| | Volumetric swelling $V_{day\ 14}/V_{day\ 1}$ |
|---|---|
| Soft-noDeg | 1.25 |
| Stiff-Deg | 1.71 |

**Supplementary Table S3:** Volumetric swelling used for calculating cell number per unit volume at day 14. N=6 hydrogels.

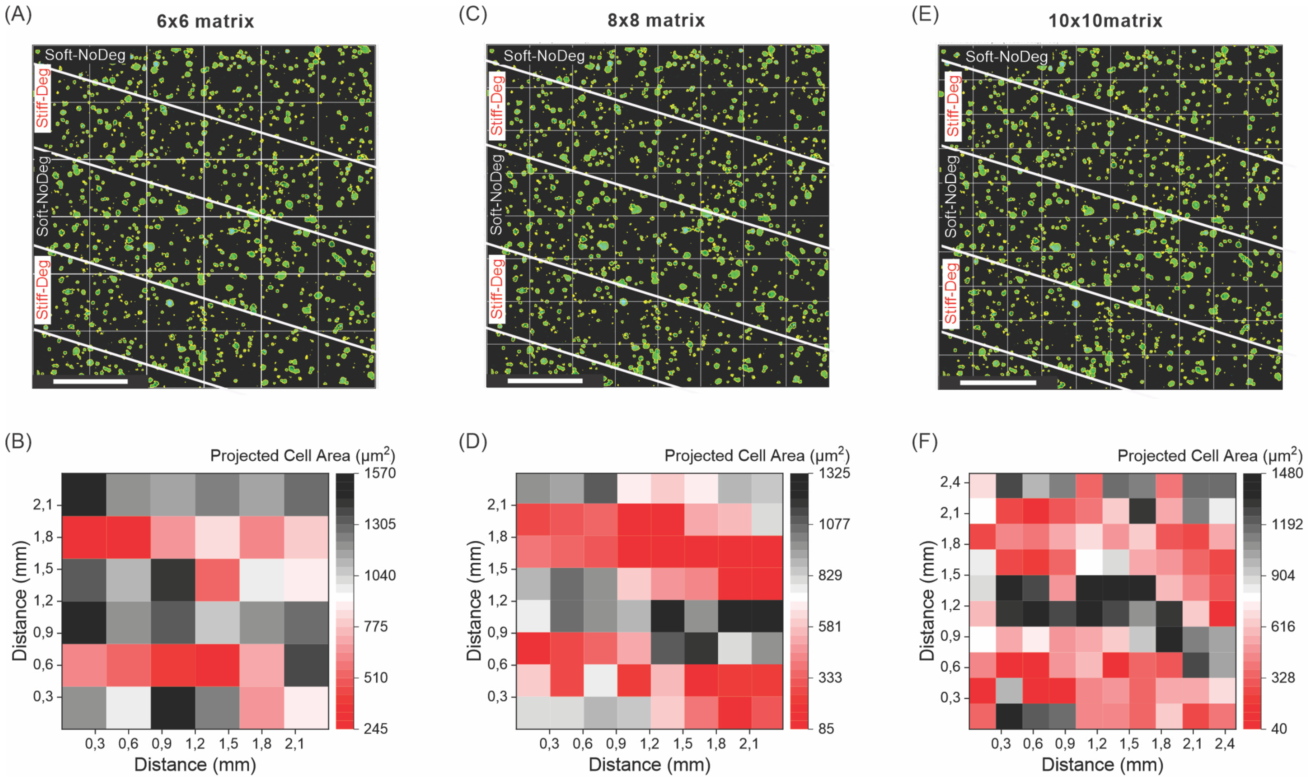


**Supplementary Figure S4:** Optimization of the binning size to visualize patterns in morphology. (A) Binning of 6x6 and (B) the respective heatmap, which shows an overestimation of the cell area and an unclear pattern; this binning size can be used for broader patterns or larger cells. (C) Binning of 8x8 and (D) the respective heatmap, which shows the optimal binning size. (E) Binning of 10x10 and (D) the respective heatmap, which has an underestimation of certain points in the pattern due to spots with no cells; this binning can be used for higher cell seeding density or smaller cells. The optimization of the image binning depends on the cell size, size of the stripes in the pattern and cell seeding density. Scale bars (A, C, E): 500µm.

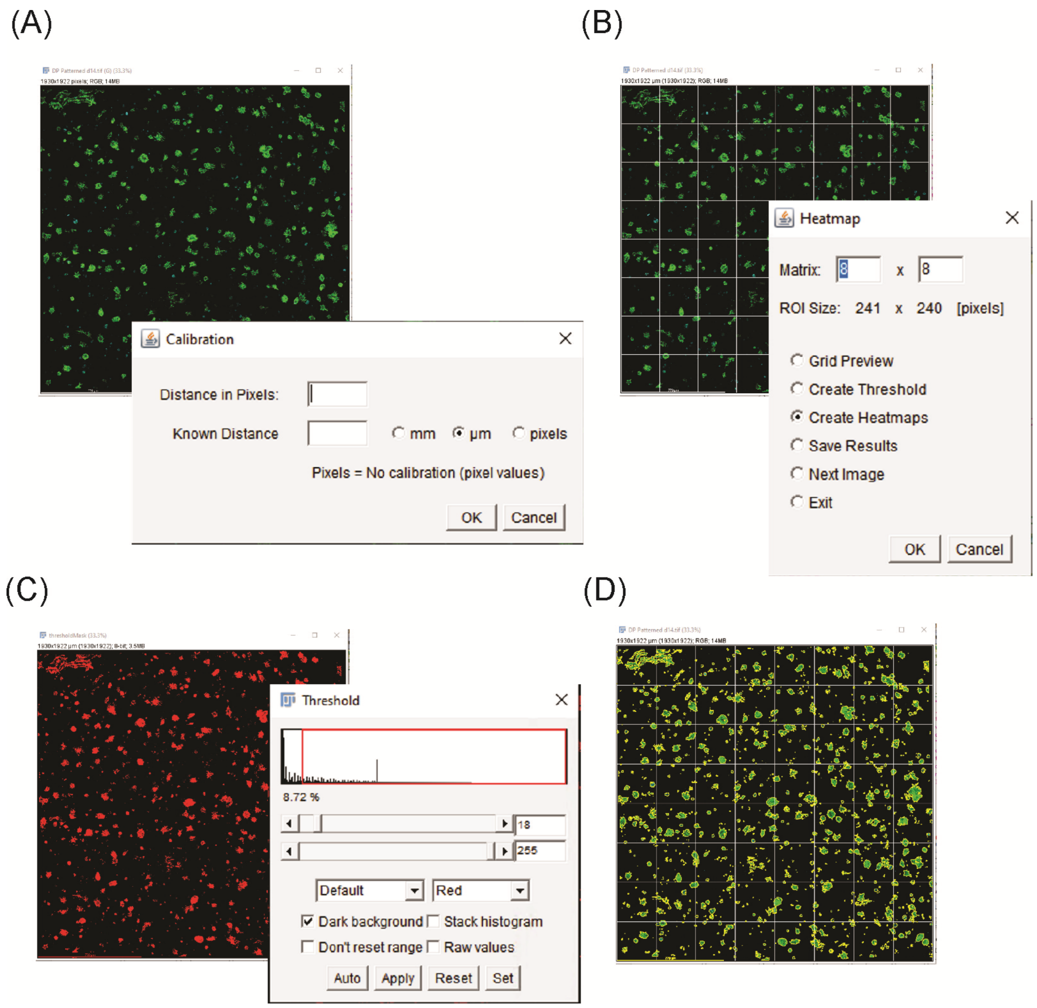


**Supplementary Figure S5:** Steps of the image-based analysis tool for cell morphology in anisotropic material analysis. (A) Setting the scale using the scale bar, (B) setting the desired binning size, (C) setting the threshold, (D) de-noising the image by setting an integer value. Finally, the heatmaps are created as data output of the software.

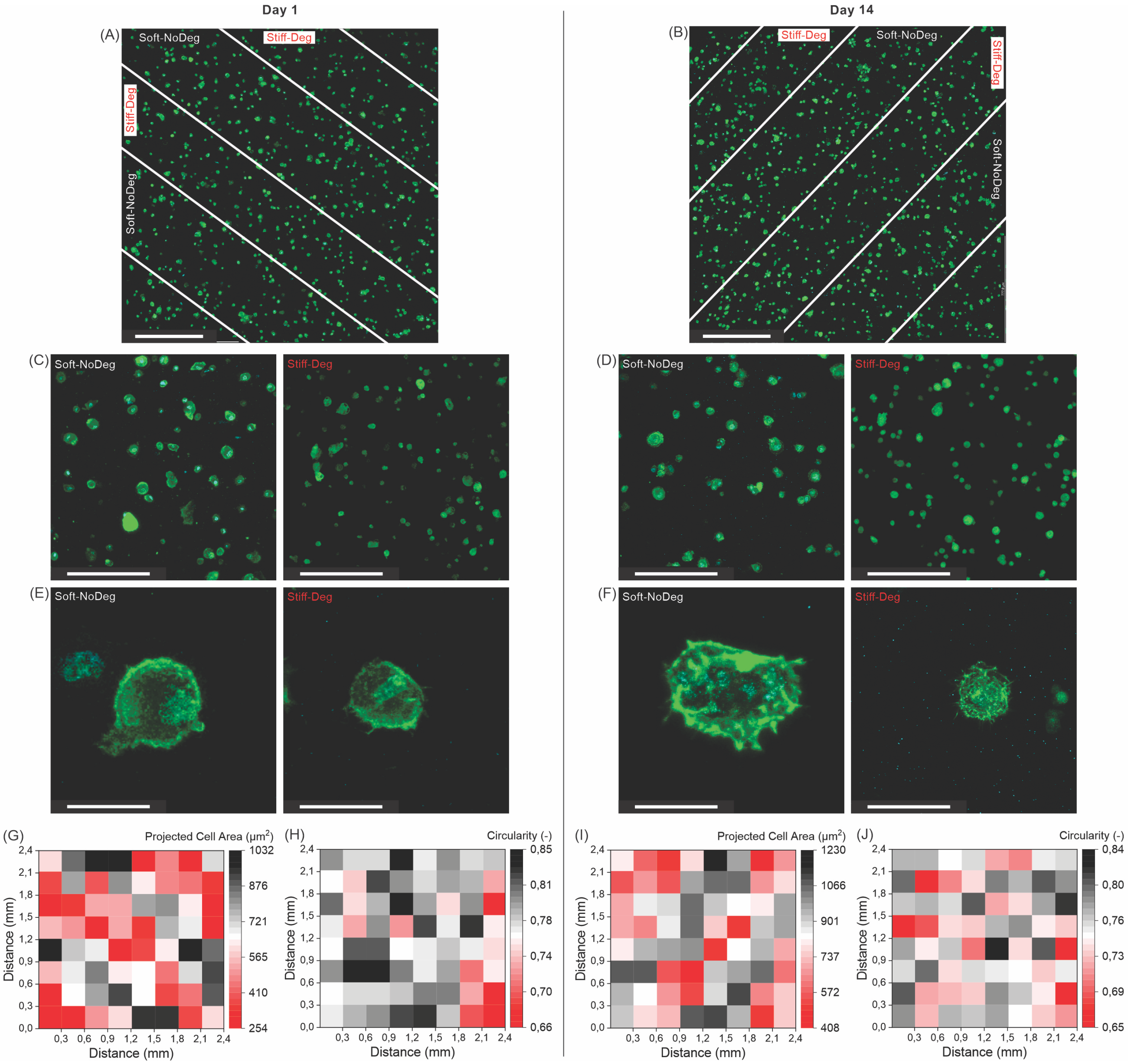


**Supplementary Figure S6:** Negative control for patterned materials using a non-degradable MMP-scramble peptide and analysis of the morphology of encapsulated cells. (A, B) Overview image with indicated pattern areas, 2x2 tile image, 10x magnification, 250 µm z-stack. (C, D) Zoom-in on the individual regions of the pattern, 25x magnification, 250 µm z-stack. (E, F) Single cell images z-stack, 40x magnification. Heat map representing (G, I) mean projected cell area ($\mu m^2$) and (H, J) circularity (-) in the overview image. Images and data from day 1 (A, C, E, G, H) and day 14 (B, D, F, I, J). Phalloidin (green)/ DAPI (cyan) staining. Scale bar: (A, B) 500 µm, (C, D) 200 µm, (E, F) 25 µm.
